# Predicting candidate genes from phenotypes, functions, and anatomical site of expression

**DOI:** 10.1101/2020.03.30.015594

**Authors:** Jun Chen, Azza Althagafi, Robert Hoehndorf

## Abstract

**Motivation:** Over the past years, many computational methods have been developed to incorporate information about phenotypes for disease gene prioritization task. These methods generally compute the similarity between a patient’s phenotypes and a database of gene-phenotype to find the most phenotypically similar match. The main limitation in these methods is their reliance on knowledge about phenotypes associated with particular genes, which is not complete in humans as well as in many model organisms such as the mouse and fish. Information about functions of gene products and anatomical site of gene expression is available for more genes and can also be related to phenotypes through ontologies and machine learning models.

**Results:** We developed a novel graph-based machine learning method for biomedical ontologies which is able to exploit axioms in ontologies and other graph-structured data. Using our machine learning method, we embed genes based on their associated phenotypes, functions of the gene products, and anatomical location of gene expression. We then develop a machine learning model to predict gene–disease associations based on the associations between genes and multiple biomedical ontologies, and this model significantly improves over state of the art methods. Furthermore, we extend phenotype-based gene prioritization methods significantly to all genes which are associated with phenotypes, functions, or site of expression.

**Availability:** Software and data are available at https://github.com/bio-ontology-research-group/DL2Vec.

## 1 INTRODUCTION

Understanding the molecular mechanisms underlying a set of abnormal phenotypes is important for diagnosis, prevention, and development of therapies. Methods to identify and study these mechanisms include observational, experimental, and computational approaches. In particular, in rare diseases, deciphering the mechanisms underlying a set of phenotypes is often limited due to small sample sizes. Computational methods that can reveal or model mechanisms in these diseases often rely on biological background knowledge.

Several computational methods have been developed to prioritize candidate genes for a particular disease or set of abnormal phenotypes (Tranchevent *et al*., 2016; Tomar *et al*., 2019; Guala and Sonnhammer, 2017; Zhang *et al*., 2018; Feng, 2017). Many such methods rely on identifying similarities between genes and suggest new candidates based on such a similarity (Gillis and Pavlidis, 2012). This similarity can be computed on several known features about a gene, including phenotype associations (Greene *et al*., 2016), distance within an interaction network (Peng *et al*., 2018), or functional similarity (Liu et al., 2018; Schlicker and Albrecht, 2007)

Phenotype-based methods have been particularly successful in finding candidate genes causing Mendelian diseases (Hoehndorf et al., 2011; Washington et al., 2009; Shefchek et al., 2019). Phenotype-based methods compare disease phenotypes to known genotype–phenotype associations and suggest candidate genes based on phenotype similarity measures (Kohler et al., 2009). While these methods are successful, their main limitation is the incomplete knowledge of phenotypes that are associated with particular genotypes. One approach to overcome this limitation is the use of phenotype associations from model organism experiments together with ontologies that integrate phenotypes across different species (Washington et al., 2009; Hoehndorf et al., 2011; Smedley et al., 2013; Bone et al., 2016; Wang et al., 2017a). Although the use of model organisms expanded the scope of prioritizing candidate genes, there is only a limited amount of information about phenotype associations available for genotypes in model organism; furthermore, genes for which there are no orthologs in model organisms cannot benefit from cross-species phenotype-based gene prioritization approaches.

One possibility to overcome the limited information on genotype– phenotype associations is the use of prediction models that predict phenotypes, and efforts such as the Computational Assessment of Function Annotation (CAFA) (Zhou et al., 2019) challenge regularly evaluate function and phenotype prediction models; while function prediction methods have increased significantly in performance and provide accurate predictions at least for some types of functions (Zhou et al., 2019), phenotype predictions still perform worse than function prediction methods (Jiang et al., 2016). Phenotypes arise from a genotype and interactions with the environment (Johannsen, 1911), and predicting the endophenotypes resulting from molecular aberrations requires the use of knowledge about molecular interactions as well as physiological interactions within and between cells, tissues, and organs.

Logical axioms, as used in many phenotype ontologies to formally characterize and standardize phenotype descriptions (Gkoutos et al., 2004; Mungall et al., 2010; Kohler et al., 2018), relate phenotypes systematically to biological functions and anatomical locations, and thereby integrate physiology, anatomy, and abnormal phenotypes within a unifying formal framework (Mungall et al., 2010; Gkoutos et al., 2018). The axioms in phenotype ontologies rely on ontologies that can be applied across different species. In particular, biological processes, functions, and cellular anatomy are described using the Gene Ontology (GO) (Ashburner et al., 2000), and anatomical sites are described using the UBERON anatomy ontology (Mungall et al., 2012); both ontologies are designed to integrate information across different species, and multiple large databases contain information that relate biological entities with classes in these ontologies. Phenotype ontologies therefore not only integrate background knowledge but can also be used to integrate data associated with different ontologies; in particular, they can be used to integrate functions of gene products, anatomical site or tissue of gene expression, and phenotypes resulting from a gene’s loss of function.

Using ontologies and the background knowledge they contain in machine learning models can significantly improve their performance (Smaili et al., 2019a). Here, we developed an ontology-based machine learning method to prioritize candidate genes based on abnormal phenotypes observed in mouse models, the normal functions of gene products, and anatomical location of gene expression. Our method combines axioms in ontologies and annotations to ontology classes. We evaluate several machine learning methods that utilize ontology axioms, and develop a novel graph-based method that overcomes several limitations of existing methods, in particular when applying machine learning to different ontologies in which classes are not related mainly through subclass axioms but rather through other types of axioms. We demonstrate that our approach improves significantly compared to established phenotype-based gene prioritization methods, and further extends the application of these methods to all genes for which either their functions or their anatomical location of expression is known.

## 2 RESULT

### 2.1 Phenotype-based prioritization of candidate genes

We developed a method based on deep learning to rank candidate causative genes given a set of abnormal phenotypes that characterize a genetically-based disease. We prioritize, or rank, genes based on three distinct types of features that can be associated with a gene: phenotypes associated with the gene’s orthologs in the mouse; the functions and cellular locations of the gene products for which the gene encodes; and the anatomical locations at which the gene is expressed. Each of these features is expressed using biomedical ontologies and we use the ontology as part of the learning problem. For this purpose, we first embed the information about genes and diseases together with the ontologies used to characterize them in a vector space and then use a supervised machine learning model to predict whether a gene is causative of a set of phenotypes or disease.

Specifically, we obtain the annotations of human genes with functions and cellular locations encoded by the Gene Ontology (GO) (Ashburner et al., 2000) from the GO Annotation database (Huntley et al., 2014), their anatomical site of expression in functional genomics experiments (Consortium et al., 2015) encoded using the UBERON anatomy ontology (Mungall et al., 2012), and the phenotypes of their mouse orthologs from the Mouse Genome Informatics (MGI) database (Smith et al., 2017) and characterized using the Mammalian Phenotype Ontology (MP) (Smith and Eppig, 2009). Furthermore, we obtain phenotype annotations of human diseases with the Human Phenotype Ontology (HPO) (Robinson et al., 2008) from the HPO database (Köhler et al., 2018). To combine the annotations using the different ontologies, we use the integrated PhenomeNET ontology (Rodríguez-García et al., 2017).

We “embed” each gene and disease, their ontology-based annotations, and the ontologies used in the annotations, in a vector space. An embedding is a function from gene or disease identifiers, and from ontologies, into 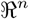 such that structural properties of the annotations and ontologies are preserved within 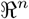. Initially, we use the Onto2Vec (Smaili et al., 2018) and OPA2Vec (Smaili et al., 2019b) methods to generate the embeddings as they have performed well in similar tasks before. We generate embeddings individually using phenotype, GO, and UBERON annotations; because these annotations are available for different numbers of genes, we also generate a set of embeddings based on the union of all genes and their annotations (i.e., for genes that have annotations from one, two, or all three datasets) as well as another set of embeddings only for genes that have annotations from all three sources.

We use a pointwise learning-to-rank model (see Materials and Methods, and Supplementary Figure 1), to prioritize gene-disease pairs based on gene–disease associations in the Online Mendelian Inheritance in Men (OMIM) database (Amberger et al., 2011). Our model is based on neural networks; given a pair of embedding vectors *G* and *D* as input, the model independently transforms the embeddings into a lower-dimensional representations using two fully-connected hidden layers, and then computes the inner product followed by a sigmoid function that outputs a value between 0 and 1, and which we use as the prediction score for an association between G and D.

We train and test our model based on 10-fold cross-validation; in each fold we split our data by the disease (and not by the gene– disease association pair) to ensure that the diseases on which we test have not been seen during training. Within each split, we use 10% of the data as the testing data used to report the final results of our model, and we use the other 90% data to train the model and tune its parameters; within these 90% of training data in each fold, we use a randomly chosen set of 90% for training and 10% for validation. We use sub-sampling of “unknown” associations between genes and diseases to generate negative associations for each disease; we sample 20 negatives for each positive association. We then use binary cross-entropy as the loss function to optimize the ranking model and use the Adam optimizer (Kingma and Ba, 2014) to train our model.

For the evaluation of our learning-to-rank model, we rank all genes for each disease based on their prediction score (within the testing set). We then use the receiver operating characteristic (ROC) curve (Fawcett, 2006) and the area under the ROC curve (ROCAUC) to evaluate how well known positive gene–disease pairs rank among all the possible pairs. Supplementary Figure 2 shows the ROC curves for our prediction model when using both Onto2Vec and OPA2Vec as embedding methods, and Table 1 summarizes the results of the cross-validation. We find that we can identify causative genes best when using phenotypes, while the predictive performance decreases when using features derived from gene functions and anatomical site of gene expression.

**Table 1.** Evaluation results for predicting gene-disease associations using embeddings generated from the Mammalian Phenotype (MP) ontology, Gene Ontology (GO), and UBERON anatomy ontology. The *intersection* represents embeddings generated jointly from all three types of ontologies and associations, limited to genes that have associations to all three ontologies, while *union* represents embeddings generated jointly from all three types of ontologies and associations, limited to genes that have associations in at least one of the three ontologies. For ROCAUC, we report the intervals obtained from cross-validation.

| Methods | Features | Gene Disease Associations without PPI |  |  |  | Gene Disease Associations with PPI |  |  |  |
| --- | --- | --- | --- | --- | --- | --- | --- | --- | --- |
|  |  | ROCAUC | Hits@1 | Hits@10 | Hits@100 | ROCAUC | Hits@1 | Hits@10 | Hits@100 |
| Onto2Vec | MP | 0.903 [0.835-0.931] | 0.012 | 0.207 | 0.355 | 0.929 [0.889-0.956] | 0.019 | 0.274 | 0.44 |
|  | GO | 0.829 [0.789-0.910] | 0.004 | 0.131 | 0.315 | 0.856 [0.832-0.917] | 0.008 | 0.226 | 0.364 |
|  | UBERON | 0.700 [0.638-0.739] | 0.003 | 0.080 | 0.145 | 0.698 [0.654-0.740] | 0.006 | 0.063 | 0.118 |
|  | Intersection | 0.918 [0.885- 0.935] | 0.012 | 0.256 | 0.442 | 0.928 [0.800-0.959] | 0.016 | 0.300 | 0.49 |
|  | Union | 0.958 [0.903-0.975] | 0.009 | 0.229 | 0.406 | 0.957 [0.929,0.968] | 0.013 | 0.258 | 0.445 |
| OPA2Vec | MP | 0.907 [0.864-0.923] | 0.019 | 0.204 | 0.372 | 0.91 [0.814-0.947] | 0.015 | 0.25 | 0.427 |
|  | GO | 0.826 [0.802- 0.862] | 0.015 | 0.194 | 0.313 | 0.841 [0.821-0.880] | 0.009 | 0.219 | 0.354 |
|  | UBERON | 0.717 [0.650- 0.784] | 0.017 | 0.127 | 0.208 | 0.727 [0.702-0.759] | 0.006 | 0.114 | 0.191 |
|  | Intersection | 0.920 [0.880- 0.936] | 0.011 | 0.256 | 0.445 | 0.928 [0.812- 0.948] | 0.020 | 0.304 | 0.468 |
|  | Union | 0.954 [0.937-0.967] | 0.008 | 0.197 | 0.378 | 0.959 [0.943-0.965] | 0.013 | 0.248 | 0.457 |
| Our model | MP | 0.957 [0.931-0.971] | 0.026 | **0.284** | 0.638 | 0.959 [0.940-0.987] | 0.037 | **0.304** | **0.650** |
|  | GO | 0.910 [0.892-0.947] | 0.023 | 0.235 | 0.579 | 0.904 [0.853-0.943] | **0.041** | 0.282 | 0.583 |
|  | UBERON | 0.842 [0.685-0.898] | 0.020 | 0.155 | 0.470 | 0.854 [0.821-0.889] | 0.019 | 0.190 | 0.515 |
|  | Intersection | 0.954 [0.928-0.977] | **0.032** | 0.252 | **0.642** | 0.953 [0.932-0.980] | 0.036 | 0.277 | 0.636 |
|  | Union | **0.978 [0.967-0.998]** | 0.024 | 0.244 | 0.635 | **0.974 [0.949-0.987]** | 0.024 | 0.230 | 0.597 |

We hypothesize that one of the reasons for the observed difference in predictive performance between the different data types is the inability of Onto2Vec and OPA2Vec to capture longer distance dependencies through which phenotypes, functions, and anatomical locations are connected within the PhenomeNET ontology. In particular, Word2Vec is equivalent to factorizing a matrix which contains the pointwise mutual information (PMI) (Church and Hanks, 1990) of words within a context window (Levy and Goldberg, 2014), and this measure is only based on directly co-occurring tokens (within the context window considered by Word2Vec). When using Onto2Vec or OPA2Vec, genes and diseases will only directly co-occur with the ontology classes used to characterize them (i.e., phenotypes, GO functions, and UBERON anatomical classes for genes, and phenotypes for diseases), as well as all their superclasses (because Onto2Vec and OPA2Vec compute the transitive closure over the subclass hierarchy and add them to the set of asserted axioms). Consequently, even if a phenotype class is defined based on an anatomical location or a function, this anatomical location or function class will not cooccur with a gene or disease that is associated with this phenotype. For example, the class *Ventricular septal defect* (`HP:0001629`) is defined as an incomplete closure of the *Interventricular septum* (`UBERON:0002094`), which in turn is constrained to be a part of the *Heart* (`UBERON:0000948`) in UBERON. When embedding genes based on their anatomical site of expression (i.e., using the UBERON ontology) and diseases based on their phenotypes, Onto2Vec and OPA2Vec will only add subclass relations as directly co-occurring tokens to use in the embedding but not the classes that are linked indirectly through axioms. We hypothesize that by incorporating these indirect associations will allow us to better utilize the background knowledge contained in the ontologies and further improve predictive performance, and we develop a novel embedding approach for ontologies that aims to improve the embedding of ontologies with many complex axioms, as well as embeddings of entities which are annotated with classes that do not stand in a subclass relation but are related through more complex axioms.

### 2.2 Embedding graph-based representations of ontologies

Our novel embedding approach is inspired by the OWL2Vec (Holter *et al*., 2019) as well as the Walking RDF & OWL (Alshahrani *et al*., 2017) methods which first convert ontologies into a graph based on syntactic patterns within the ontology axioms, and then apply a knowledge graph embedding (Wang *et al*., 2017b) on the resulting graph. However, our method extends OWL2Vec and Walking RDF & OWL to incorporate more complex forms of axioms into the generated graph so that the complexity of the axioms in a cross-species phenotype ontology such as PhenomeNET (Rodríguez-García *et al*., 2017) can be utilized.

We have defined a transformation function that is used to convert ontology axioms in the Web Ontology Language (OWL) (Grau *et al*., 2008) format into a graph. The transformation function considers logical operators as well as quantifiers, and converts them into edges (or subject–predicate–object triples) of a graph. Our transformation considers axioms pertaining to classes (the ontology, or TBox). Associations between a gene *G* and an associated ontology class *C* can be modelled in OWL as an axiom `G SubClassOf: has-function some C` (or using some other relation instead of `has-function`) and, as a consequence, a direct edge between G and C will be created through our algorithm. We convert all axioms from the PhenomeNET ontology, and the annotations of gene and disease entities with their ontology classes, into a graph representation using the transformation function in Table 2. After generating the graph, we apply iterated random walks starting at nodes of the graph to generate a corpus, and use Word2Vec (Mikolov et al., 2013) to generate embeddings for nodes and edge labels based on this corpus.

**Table 2.** The transformation rules to convert ontology axioms into a graph. *Q* represents an arbitrary quantifier or cardinality restriction.

| Condition 1 | Condition 2 | Triple(s) |
| --- | --- | --- |
| $A \sqsubseteq QR_0 \dots QR_mD$<br>$A \equiv QR_0 \dots QR_mD$ | $D \equiv B_1 \sqcup \dots \sqcup B_n \parallel B_1 \sqcap \dots \sqcap B_n$ | $\langle A, (R_0 \dots R_m), B_i \rangle$ for $i \in 1 \dots n$ |
| $A \sqsubseteq B$ | | $\langle A, SubClassOf, B \rangle$ |
| $A \equiv B$ | | $\langle A, EquivalentTo, B \rangle$ |

We repeat our supervised training process using our novel embedding method. The results are summarized in Table 1 and ROC curves for this task shown in Supplementary Figure 2. While the performance in predicting gene–disease associations using only phenotype annotations is comparable to the predictive performance observed when using Onto2Vec and OPA2Vec, we observe a significant improvement when using features encoded using GO (*p* = 2.02 × 10^-125^, Mann-Whitney U test) and UBERON (*p* = 2.77 × 10^-152^, Mann-Whitney U test), indicating that our approach can better capture relations between classes that are related through complex axioms instead of only subclass axioms.

### 2.3 Adding network information

Since our embedding approach is based on graphs and random walks, it can naturally accommodate other graph-structured information in addition to the graph generated from the ontology axioms. There are many biological networks that also relevant to understanding gene–disease associations (Alanis-Lobato et al., 2016; van Dam et al., 2018; Al-Harazi et al., 2016), in particular interaction networks. To determine whether our method is able to utilize this information, we conduct another experiment in which we add functional interactions between proteins obtained from the STRING database (Szklarczyk et al., 2019) to the knowledge base. We add the interactions to our graph as direct `interacts-with` edges between genes, and then we repeat our workflow and predict associations between genes and diseases based on the new embeddings.

The results of this experiment are shown in Table 1 and Supplementary Figure 3, which shows the overall performance obtained from our method using network information and its comparison with embeddings based on Onto2Vec and OPA2Vec. We find that our workflow results in the best-performing model when using our method to generate embeddings in particular when comparing the embeddings generated using ontologies of different domains, such as when comparing diseases (characterized with phenotypes) and genes characterized by their function or anatomical site of expression; and that our method is effectively able to incorporate additional network information.

## 3 DISCUSSION

We designed a novel method to prioritize candidate genes given a set of abnormal phenotypes associated with a genetically-based disease; our method uses information about genes obtained from animal model phenotypes, the functions of gene products, the anatomical location of gene expression, and interaction networks, as well as a large amount of background knowledge contained in biomedical ontologies. Our method improves over other phenotype-based methods in several ways.

First, we use a pointwise learning-to-rank machine learning model which improves the predictive performance when evaluated using gene–disease associations from the Online Mendelian Inheritance in Men (OMIM) (Hamosh et al., 2005) database; our model is designed to directly learn the similarities between two embeddings and results in improved predictive performance when compared to other models (Smaili et al., 2019b, 2018) used to predict gene–disease associations based on embeddings.

Second, we developed a novel method to exploit complex axioms by converting them into a graph and relying on graph embeddings; we show that this approach improves performance significantly when embedding multiple ontologies that are only linked through complex axioms. This advance is particularly important in ontologies that are heavily formalized using OWL and that are interlinked, such as the ontologies in the collaborative OBO Foundry effort (Smith et al., 2007).

Third, our method is, to the best of our knowledge, the first that prioritizes candidate genes for a set of abnormal phenotypes using a combination of gene expression, function, network, phenotype data, and ontologies. While many of these features have been used individually to rank candidate genes (Singleton et al., 2014; Schlicker and Albrecht, 2007; Smedley et al., 2013), they have not been combined in a single model. Crucially, our method combines this information on two distinct levels: first, the different annotations (phenotype, function, expression) are combined on the level of a gene or gene product (which we do not distinguish), so that a single entity (the gene and its products) is associated with all three types of information; second, we also utilize the links between ontologies directly. The links between the classes in ontologies allow us to establish new relations between the different features associated with genes, and these features are not accessible without utilizing the ontology axioms.

Our method still has several limitations. Our conversion from OWL into a graph does not consider all OWL axioms, and the conversion also treats different types of restrictions and axiom types identically although their semantics is different. In the future, we plan to extend the method to convert any OWL axioms into a graph representation, relying, for example, on relational patterns defined in the OBO Relation Ontology (Smith *et al*., 2007), and also rely on inferred axioms for generating the graph such as implemented in the Onto2Graph method (Rodríguez-García and Hoehndorf, 2018).

Another major limitation of our approach is that it is inherently transductive and not inductive. In particular, the diseases with their phenotype associations must be known in our workflow before generating embeddings and training our prediction model, and it is not straightforward to apply the approach to a new set of phenotypes (such as the phenotypes observed in an individual). This limitation is shared by many graph embedding and knowledge graph embedding approaches (Wang *et al*., 2017b). In the future, we may overcome this limitation through the use of inductive methods for learning on knowledge graphs, such as graph neural networks (Scarselli *et al*., 2008; Kipf and Welling, 2016). Extending our approach to an inductive setting will also allow use to combine our approach with methods to find pathogenic causative variants based on observed phenotypes and next generation sequencing data (Metzker, 2010).

## 4 CONCLUSIONS

We developed a method for prioritizing candidate genes given a set of phenotypes associated with a disease. Our method can utilize different types of features characterized through ontologies, and significantly improves the phenotype–based prediction of disease genes. While previous phenotype–based gene prioritization methods are only applicable when phenotype associations are known for genes, our method can be applied to a much larger number of genes for which either functions, sites of expression, phenotypes, or interactions with other genes are known.

## 5 MATERIALS AND METHODS

### 5.1 Ontology and annotation resources

We downloaded Gene Ontology (GO) (Ashburner *et al*., 2000) annotations of 18,495 human gene products (495,719 annotations in total) from the Gene Ontology website on 2020-03-20. We excluded the GO annotations where the evidence code indicated that the annotation was inferred from electronic annotation (IEA) or for which no biological data is available (ND). We map the UniProt accessions to Entrez gene identifiers using the mappings provided by the Entrez database (Maglott *et al*., 2010), downloaded on 2020-03-20, and we obtained 17,786 Entrez genes where the gene product has GO annotations.

We obtained phenotype annotations for 13,529 mouse genes, including 228,214 associations between genes and Mammalian Phenotype Ontology (Smith and Eppig, 2009) classes, from the file `MGI_GenePheno.rpt` available at the Mouse Genome Informatics (MGI) database (Smith *et al*., 2017). Phenotype associations were downloaded on 2020-03-20. We map each mouse gene to their human ortholog using the file `HMD_HumanPhenotype.rpt` available at the MGI database, resulting in 9,879 human genes where the mouse ortholog has phenotype associations.

We further downloaded the Tissue Expression Profiles (GTEx) dataset (Consortium *et al*., 2015) from the Gene Expression Atlas (Papatheodorou *et al*., 2019) on 2020-03-20. GTEx characterizes gene expression across 53 tissues. We map the Ensembl protein identifiers to Entrez gene identifiers using the mapping provided by the Entrez database (Maglott *et al*., 2010). We set a threshold for whether a gene is expressed or not in a tissue by setting a cutoff of 4.0 transcripts per million (TPM); this threshold is determined experimentally (see Supplementary Figure 4). Finally we obtained 20,538 Entrez genes which have expression above this threshold in one or more tissue. We map each tissue to the Uberon Anatomy Ontology (Mungall *et al*., 2012), downloaded from the AberOWL ontology repository on 2020-03-20. We exclude the expression in *EBV-transformed lymphocyte* and *transformed skin fibroblast,* since these two tissues are not available in the Uberon ontology.

The PhenomeNET Ontology (Rodríguez-García *et al*., 2017) is a crossspecies ontology which integrates multiple species-specific phenotype ontologies as well as related ontologies such as the Gene Ontology and the Uberon Anatomy Ontology. We downloaded the PhenomeNET ontologyfrom the AberOWL ontology repository on March 20, 2020.

### 5.2 Evaluation datasets

We obtain associations between 2,542 human diseases and 2,885 genes from the file `MGI_DO.rpt` available at the MGI database, downloaded on 2020-03-20; the dataset contains 4,051 gene-disease associations in total, where diseases are represented using their OMIM identifier (Amberger *et al*., 2011).

As our gene–phenotype associations are to mouse genes (resulting from a loss of function of that gene) while our evaluation set uses human gene identifiers, we need to identify human orthologs of the mouse genes. We identify the mouse orthologs of human genes, and human orthologs of mouse genes, using the file **HMD_HumanPhenotype.rpt** at the MGI database, downloaded on 2020-03-20. Table 3 summarizes our training and evaluation data.

**Table 3.** Training and evaluation data used in our method. We list the annotations with each ontology, MP, GO, and UBERON, as well as the number of genes annotated with them and the number of diseases that these genes are associated with. *Intersection* represents the genes (and their associations) that have associations with all three ontologies, while *Union* represents the number of genes that have associations in one, two, or all ontologies.

| Ontology | Genes | Diseases | Associations |
| --- | --- | --- | --- |
| MP | 10,951 | 1,784 | 3,888 |
| GO | 17,786 | 1,784 | 3,883 |
| UBERON | 20,538 | 1,691 | 3,687 |
| Intersection | 9,886 | 1,687 | 3,655 |
| Union | 22,707 | 1,787 | 3,905 |

We use functional interactions between proteins obtained from the STRING database (Szklarczyk *et al*., 2019) on March 01, 2020. The interaction dataset contains 19,355 proteins and 11,759,455 edges between them. We mapped the proteins to the UniProt database and filter out those entries that did not map to the UniProt database. Further, STRING provides a confidence score for an interaction and we only keep interactions with a confidence of at least 700. The remaining interaction network consists of 17,178 proteins with 840,672 interactions.

### 5.3 Embedding methods

Onto2Vec (Smaili *et al*., 2018) is a method to learn the semantic embedding representations of biological entities and by extracting features from ontology-based annotations, axioms and ontology structures. It directly utilizes the axiom features and also indirectly infer new logical axiom features by applying the HermiT OWL reasoner (Shearer *et al*., 2008). Onto2Vec collects data and axioms as “sentences” and uses a skip–gram model to learn the vector representation for each word. OPA2Vec (Smaili *et al*., 2019b) is an extension of Onto2Vec which includes the annotation axioms in ontologies and uses transfer learning to assign them a semantics.

A random walk of length *k* on a graph *G* = (*V, E*) is a sequence of nodes and edges *n*_1_, *e*_1_,…,*e*_*k*-1,n_k__ such that for all *i*, 1 ≤ *i* < *k*, *e_i_* ≡ (*n_i_*, *n*_*i+i*_) ∈ *E* and *n*_*i*+1_ was chosen randomly from all neighbors of *n_i_*. Here, we also include edge labels in the walk. For the purpose of selecting neighboring nodes, we treat the graph as undirected. We generate 80 walks from each node, and stop the walks after 20 steps.

We implement our embedding algorithm in a software called DL2Vec and make the source code, together with our experiments, freely available under the GNU General Public License version 3.

### 5.4 Word2Vec model

Word2Vec (Mikolov et al., 2013) is a language model for learning vector representations of words based on co-occurrence within a context window. We use the skipgram model of Word2Vec (Mikolov et al., 2013). Given a sentence with *N* words, the skipgram model reads the sentence with a window kernel size c and maximizes the co-occurrence probability of words that appear in the same window.

We apply the SkipGram algorithm on our node and edge sequence corpus, which is generated by a random walk on the heterogeneous graph. We set the skipgram parameters to a window size of 10, and min_count value to 1. The training process iterates 20 times, and it outputs a 200 dimensional embedding for each entity.

### 5.5 Pointwise learning-to-rank prediction model

We use a pointwise learning-to-rank model to predict associations between genes and diseases. The model takes two vectors *V*_1_ and *V*_2_ as input for two independent neural networks and *ν*_2_. We then calculate the inner product of *ν*_1_(*V*_1_) and *ν*_2_(*V*_2_) and use a sigmoid function to obtain a similarity score between *V*_1_ and *V*_2_. We train this model using binary cross entropy as loss function. Each neural network *ν*_1_ and *ν*_2_ consists of two hidden layers, the first with 256 neurons and the second with 50. We use 20% dropout (Srivastava et al., 2014) after each layer, followed by a LeakyReLU (Xu et al., 2015) as the activation function. The model parameters are optimized using the Adam (Kingma and Ba, 2014) optimizer.

### 5.6 Evaluation

We use the Receiving Operating Characteristic (ROC) curve (Fawcett, 2006) to assess the performance of our classification model. The ROC curve is a plot of the true positive rate as a function of the false positive rate. To compute true positive and false positive rate, we rank all genes for each disease, and compute the average true and false positive rates at each rank. We then generate the ROC curve, and compute the area under the ROC curve, as the averages across all diseases. We also report the recall at rank n (Hits@n).

We compute differences in the area under the ROC curve using the nonparametric Mann Whitney U test (Nachar et al., 2008). For the test, we test the significance of ranking true positive associations differently between two prediction models. We consider differences as significant if *p* < 0.05. In order to compare the performance of the embeddings generated from phenotypes (MP), gene expression (UBERON), and biological functions (GO) directly, we focus on genes which have annotations to all three ontologies as evaluation set; the number of genes that have annotations in all three ontologies is 9,886.

