## Supplementary for "Predicting candidate genes from phenotypes, functions, and anatomical site of expression"

#### 1 Selection of gene expression thresholds

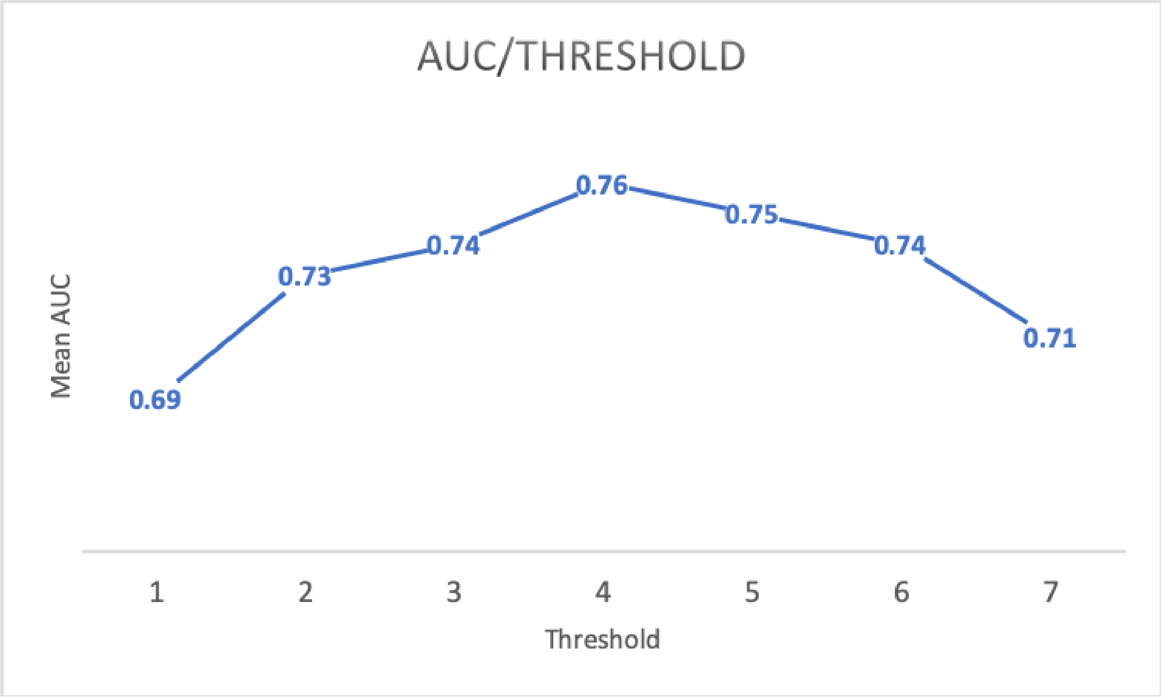


Figure 1: Onto2Vec AUC performance in 10-fold crossvalidation using different thresholds for gene expression in tissues.

#### 2 Ranking model

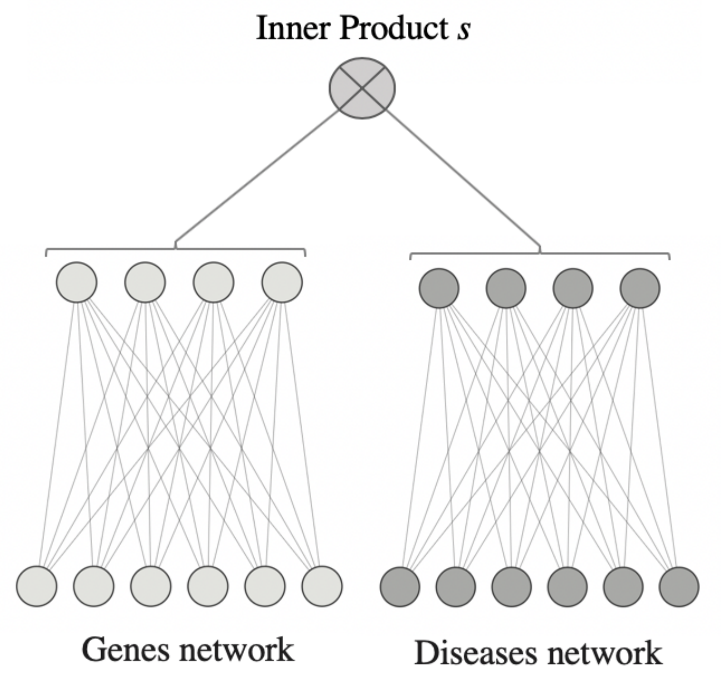


Figure 2: Ranking model with pointwise loss. We use this model to predict whether there should be a relation between the gene and disease (both of which are represented as "embeddings").

### 3 ROC curves for predicting gene–disease associations

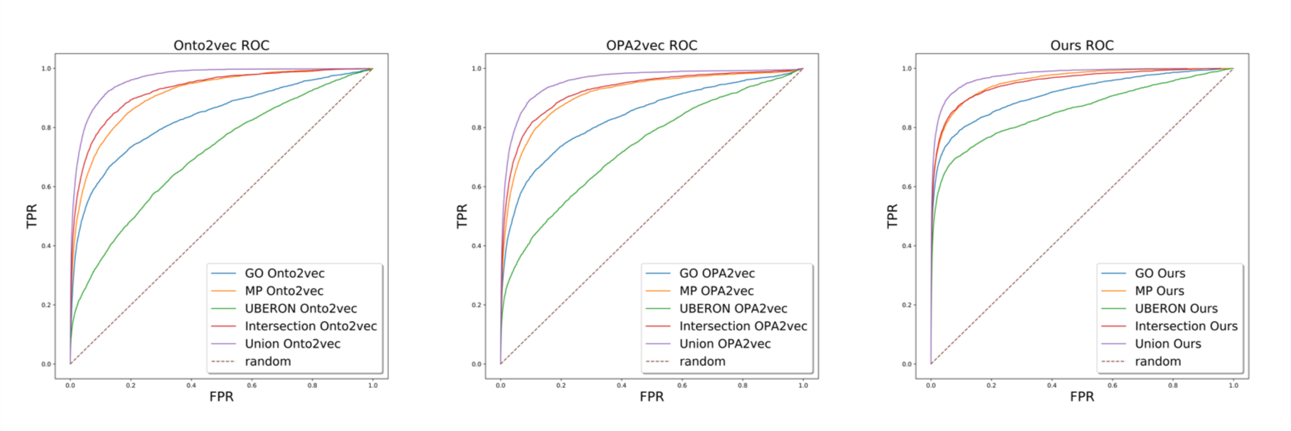


Figure 3: ROC curves for predicting gene–disease associations based on Onto2Vec, OPA2Vec and our method.

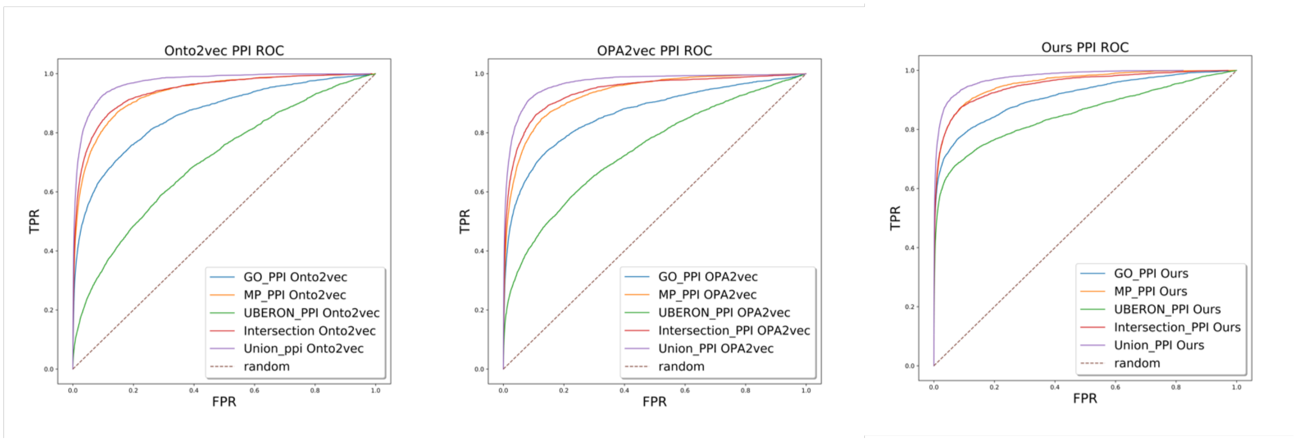


Figure 4: ROC curves for predicting gene–disease associations based on Onto2Vec, OPA2Vec and our method when including interaction data.
